# Optimized *k*-mer search across millions of bacterial genomes on laptops

**DOI:** 10.1101/2025.11.23.690050

**Authors:** Francesca Brunetti, Karel Břinda

## Abstract

Comprehensive bacterial collections have reached millions of genomes, opening new opportunities for point-of-care diagnostics and epidemiological surveillance. However, local real-time search over such collections on commodity hardware remains difficult. Currently, only LexicMap and Phylign enable local search and alignment at such a scale; among them, only Phylign is designed to run on laptops, via a subindex approach informed by phylogenetic compression. However, Phylign’s performance deteriorates on long and divergent queries because it uses COBS as a *k*-mer-based prefilter before alignment with Minimap2. Meanwhile, recent *k*-mer indexes such as Fulgor and Themisto have emerged, but there is no practical methodology for selecting, combining, and parameterizing them for phylogenetically partitioned million-genome search under constraints.

Here, we develop an end-to-end methodology for *k*-mer matching in phylogenetically compressed bacterial collections. We formalize a matching strategy defined by matching mode, query type, and reference characteristics, and use this to shortlist candidate indexes and benchmark them under space–time trade-offs. As a case study, we address plasmid search over AllTheBacteria, compare multiple index types, and identify configurations optimizing the Pareto frontier of space and speed. Guided by these results, we implement a phylogenetically compressed variant of Fulgor, integrate it into Phylign, and obtain Phylign-Fulgor, a laptop-ready pipeline for million-genome search. On the 661k collection, Phylign-Fulgor makes the prefiltering step ∼4× faster than Phylign-COBS at the cost of a 1.2× larger index. On AllTheBacteria, its *k*-mer filter is 20×–300× faster in real time than LexicMap’s alignment-based search and uses ∼20× smaller disk space. The full Phylign-Fulgor workflow including Minimap2 alignments is slower than LexicMap for a single plasmid but competitive or faster for batched plasmid queries. Phylign-Fulgor has comparable matching sensitivity to LexicMap, is less sensitive at the alignment level, but always stays within a laptop RAM budget (∼5×–20× lower memory than LexicMap).

## INTRODUCTION

Advances in DNA sequencing technologies have driven an exponential expansion of public microbial genome archives [1–5]. The European Nucleotide Archive (ENA) alone grew from 661,405 curated bacterial genomes of the “661k” Illumina snapshot [3] as of 2018 to over 2.4 million by August 2024, as captured in AllTheBacteria [4]. Modern collections of assembled bacterial genomes such as AllTheBacteria [4], NCBI’s bacterial assemblies, and GTDB [12] now contain thousands of species and millions of genomes. These databases open up qualitatively new possibilities for point-of-care applications, including rapid AMR diagnostics [8], plasmid and mobile-element tracing [9, 10], and large-scale epidemiological surveillance [11, 12]. Yet such applications ultimately rely on fast and sensitive local search and alignments across these million-genome collections on commodity hardware.

Traditional alignment tools such as BLAST [13] remain highly sensitive but become impractical at this scale: they typically require substantial memory and compute, which makes real-time search over millions of genomes infeasible. A common alternative is to use *k*-mer indexes [14], which reduce genomes to *k*-mer sets and search based on determining which query *k-*mers belong to which reference sets. However, even with *k*-mer indexes, real-time local search across million-genome collections remains challenging on laptops. State-of-the-art *k*-mer indexes either do not scale to millions of genomes [15, 16] or assume access to powerful clusters or servers [17, 18]. Other large-scale search frameworks, such as BIGSI-like systems including BIGSI [17], Metagraph [19], Logan Search [20], or general tools such as MMseqs2 [21] and RopeBWT3 [22], also rely on substantial computational infrastructure. LexicMap provides fast alignment against multi-million-genome bacterial collections but targets server-class hardware and multi-terabyte disk space [22].

To our knowledge, only LexicMap [23] and Phylign [5] currently provide sensitive local search and alignment at the scale of million assembled genomes. LexicMap enables fast alignment across AllTheBacteria and similar-scale collections but uses multi-terabyte indexes and large memory, which makes it unsuitable for portable devices. Phylign, in contrast, is designed for laptops: it uses phylogenetic compression [5] to partition genomes into batches of closely related, phylogenetically ordered genomes. Phylign builds separate COBS [23] *k*-mer indexes per batch and searches them independently before aggregating results, optionally followed by targeted Minimap2 [24] alignment for best-scoring matches. This subindex paradigm enabled, for the first time, gene- and plasmid-level searches against the 661k collection [3] in hours on a laptop.

Over the last few years, more advanced *k*-mer indexes such as Fulgor [15], Themisto [16], and Metagraph [27] have emerged. These tools offer substantially better scaling than COBS and in principle could be used as phylogenetic subindexes to accelerate search on larger collections and improve sensitivity on long or divergent queries. In practice, however, integrating them into subindex-based workflows is non-trivial. It requires a principled way to map a biological question to a concrete *k*-mer matching strategy: which *k*-mers must be matched, which matching modes are acceptable, and which index features (for instance, thresholding schemes) are admissible under the subindex paradigm.

Despite the broad ecosystem of *k*-mer indexes now available [15–17, 23, 25–40], there is still no practical end-to-end framework that (1) maps concrete biological questions to implementable *k*-mer matching strategies, (2) selects compatible indexes and configurations under (laptop) hardware constraints, and (3) evaluates disk–memory–time trade-offs when large collections must be partitioned into multiple subindexes, as in Phylign. This gap – both methodological and implementation-related – makes it difficult to move beyond tool-by-tool benchmarks and to design laptop-centric workflows for millions of genomes in realistic point-of-care use cases, such as plasmid surveillance.

Here, we make four contributions. First, we introduce a formal framework for defining *k*-mer matching strategies under the subindex paradigm, based on three elements – matching mode, query type, and reference characteristics – which together determine admissible *k*-mer indexes and configurations. Second, we develop a procedure based on Pareto-optimization for selecting *k*-mer indexes across phylogenetic batches, jointly optimizing disk usage and search time at the collection scale. Third, we apply this framework to plasmid search across AllTheBacteria, benchmarking COBS against three modern *k*-mer indexes – Fulgor, Themisto, and Metagraph – and identify Fulgor as the best trade-off index. Fourth, we integrate a phylogenetically compressed variant of Fulgor into Phylign, obtaining Phylign-Fulgor and evaluating it on the 661k and AllTheBacteria collections under realistic laptop and server scenarios.

## RESULTS

### Methodology for optimizing local *k*-mer-based search

We developed a methodology for optimizing *k*-mer search on laptops (**Fig. 1**). Given a biological question (**Fig. 1a**), we assume it can be answered with *k*-mers and formalized as a pseudo-algorithm containing a *k*-mer matching step. Its formulation dictates a *matching strategy* defined by three components: *matching mode, reference collection characteristics*, and *query type* (**Note S1**).

**Fig. 1.**
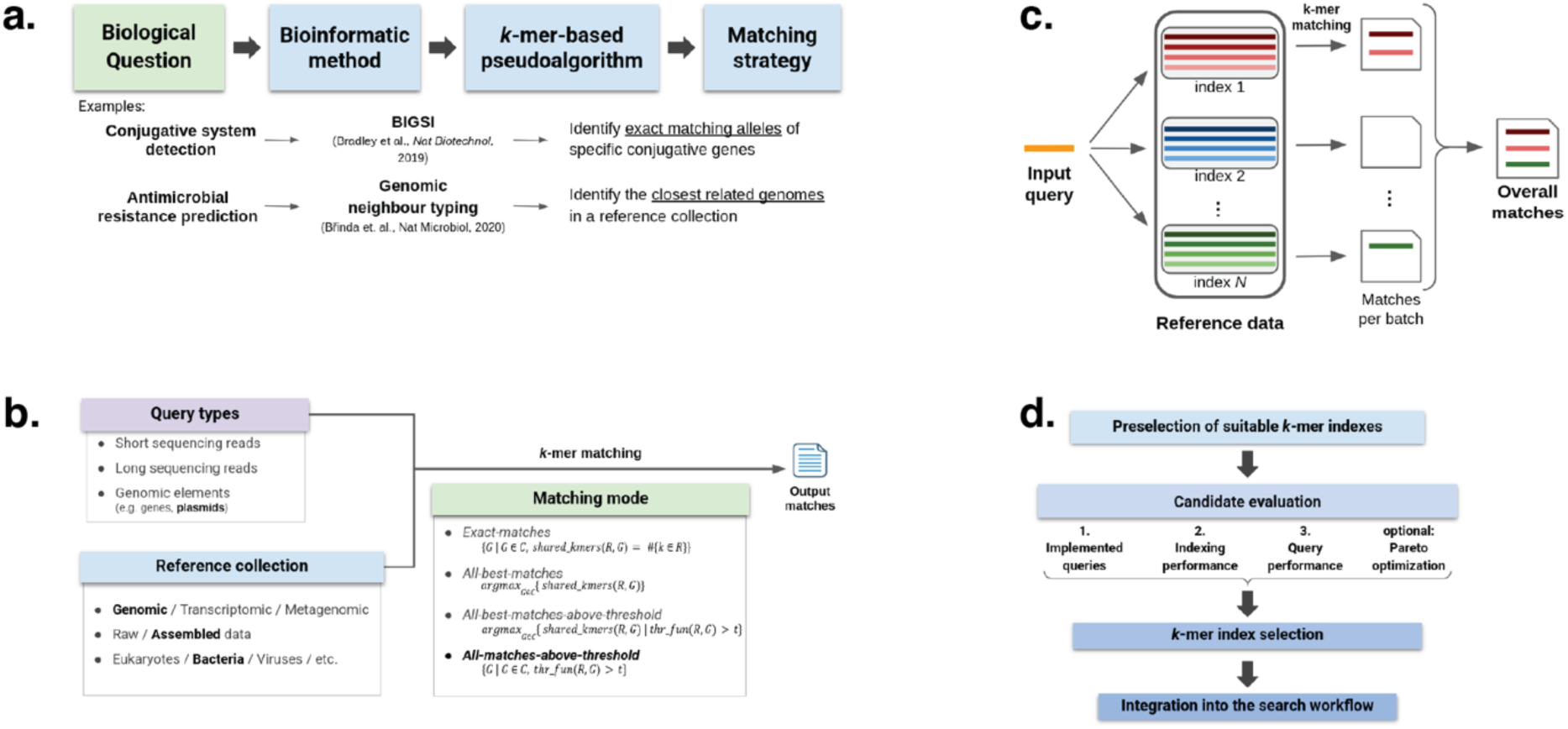
Overview of the *k*-mer selection methodology for large-scale microbial search. **a)** From biological questions to *k*-mer matching strategies. A domain question (e.g., detecting conjugative systems or predicting AMR) is reformulated as a *k-*mer-based pseudo-algorithm, which fixes the required *k*-mer matching strategy and output. **b) Matching-strategy design space.** Query type (short/long reads, assemblies, genetic elements), reference collection (genomic/metagenomic/pangenomic; raw/assembled), and matching mode (exact search / best-match / thresholded) jointly determine the admissible matching strategy. **c) Subindex search paradigm.** Large reference collections are partitioned into subindexes that are queried independently; their match sets are then aggregated into global results. **d) *k*-mer index selection process.** Candidate indexes are preselected based on technical and functional properties, benchmarked on relevant query types under disk–memory–time constraints and, optionally optimized via Pareto analysis. The final index or combination of indexes is integrated into the workflow.

Among the many available *k*-mer indexes [15–17, 23, 25–39], only a subset is relevant to any specific problem (**Fig. 1b**). Matching modes determine how *k*-mers are compared, query type (short/long reads, genes, plasmids, etc.) affects performance through length, quality, and divergence; and reference characteristics such as phylogenetic structure influence index size and search complexity – for example, clonal collections compress well and yield small, fast indexes, whereas heterogeneous ones inflate size and query time. Specific combinations of these elements define the matching strategy for a given biological question.

However, not every *k*-mer index can implement a chosen matching strategy under the *subindex paradigm*. To maintain searchability as collections grow faster than storage and memory, we rely on this paradigm as implemented in Phylign [5]: genomes are grouped into phylogenetically informed, highly compressible *batches* (**Fig. 1c**), each batch indexed and queried independently before aggregation. Complemented with phylogenetic structure [5], this enabled alignment across the full 661k collection on laptops [5]. This paradigm excludes some index designs. For instance, Themisto’s intersection regime evaluates thresholds only on k-mers present somewhere in the full index; k-mers absent in one batch but present in another break aggregation. The same limitation applies to Fulgor’s full-intersection method. The final step is to identify the most suitable *k*-mer index from a shortlist of *candidates* via benchmarking (**Fig. 1d, Note S2**).

### Case study: plasmid search over AllTheBacteria on laptops

Let us consider a specific use case: matching one or multiple plasmids on a laptop against the 661k/ATB collections. Plasmids are extrachromosomal DNA elements that facilitate horizontal gene transfer between bacterial isolates of the same or different species [42]. They can greatly vary in length and content, and their intrinsic heterogeneity undermines both *k*-mer-based filtering and alignment sensitivity at large scale. This mosaic architecture routinely breaks standard aligners – conserved regions inflate hit lists with non-informative matches, whereas divergent regions drop below sensitivity.

Here, we specifically consider the search of the EBI database of 2,826 plasmids, whose search was previously benchmarked in [5, 17], plus one additional plasmid from [22] used as a key benchmark dataset therein. Following previous work on plasmid matching using *k*-mers [5, 17], we adopt the *all-matches-above-threshold* matching mode between plasmid sequences and a reference collection of bacterial assemblies.

#### Technical and functional constraints on candidate k-mer indexes

We selected four indexes to benchmark for plasmid search within the subindex paradigm: COBS [23] as the current solution in Phylign, and Fulgor [15, 25], Themisto [16], and Metagraph [18] as potential replacements (**Tab. 1, Tab. S1**). To ensure sensitivity and accuracy, we considered only exact indexes.

**Table 1.** Comparison of functional capabilities of COBS, Fulgor, Themisto, and Metagraph. Panels report support for exact indexing and subindex compatibility (**a**) and for individual matching modes (**b**). Functionality is classified as supported (S), partially supported or requiring minor code changes (PS), or not supported (NS). For details, see **Note S3**.

|  | COBS [23] | Fulgor [15, 25] | Themisto [16] | Metagraph [18] |
| --- | --- | --- | --- | --- |
| **a. Index properties** |  |  |  |  |
| exact indexing | NS | S | S | S |
| subindex compatibility | S | PS | PS | S |
| **b. Matching modes** |  |  |  |  |
| exact-matches | S | NS | NS | S |
| best-matches | S | NS | NS | S |
| best-matches-above-threshold | S | NS | NS | S |
| all-matches-above-threshold | S | PS | PS | S |

S: Supported; PS: Partially supported; NS: Not supported

First, we assessed their compatibility with the subindex paradigm (**Tab. 1a)**. Under this paradigm, the reference collection is partitioned into multiple batches, each batch is indexed and queried independently, and the batch-wise outputs are then aggregated into a global result.

Formally, let C be a reference collection, Q a query, and M(C, Q) a well-defined *k*-mer-based similarity measure between C and Q that we wish to obtain. Let Ic denote an index built on the full collection C, and {I_j_}_j=1…n_ a family of subindexes forming a disjoint partition of C. Let out_Ic_(Q) be the output of searching Q against Ic, and out_Ij_(Q) the output against I_j_. We say that an index family is *compatible with the subindex paradigm* if there exists an aggregation function Comb such that, for all Q:

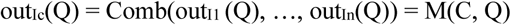

This rules out index designs whose semantics depend on the global *k*-mers presence rather than per subindex behavior.

Metagraph and COBS satisfy subindex compatibility directly through their threshold-based modes (termed *exact k-mer matching* and *approximate pattern matching* by their respective authors), returning all references whose similarity to the query exceeds a user-defined threshold computed over the full set of query *k*-mers. Themisto and Fulgor also implement thresholded search, but their default mode normalizes similarity positive *k*-mers only. This makes the denominator batch-dependent and breaks aggregation: each batch computes a different normalizing constant. To avoid this, both tools must operate in their full-*k*-mer regime and output explicit similarity values, which requires the “--include-unknown-kmers” parameter in Themisto and minor code changes in Fulgor. Under these conditions, the all-matches-above-threshold mode becomes implementable for all four tools (**Tab. 1b**).

#### Disk–-time trade-offs differ between index designs

Next, we sought to understand the time-memory trade-offs of the four selected indexes when used within the subindex paradigm. For both Fulgor and Themisto, we considered their two color-annotation variants: Fulgor/meta-colored Fulgor (we will refer to it as Fulgor and Fulgor-m, respectively) and Themisto-hybrid/Themisto-roaring bitmap (Themisto-h and Themisto-r, respectively). For Metagraph, we considered only its row-diff relaxed BRWT variant, suggested by the authors as the best trade-off between compression and query performance. As the subindex paradigm enables storing subindexes in a compressed form and decompressing them on the fly, like implemented in Phylign with COBS, we primarily considered for each index both its variant compressed by XZ. Nevertheless, as *k*-mer indexes have substantially improved recently in internal compression [15, 16, 18], we also evaluated the index size without XZ compression.

As a reference database, we used the 661k collection, as extensive calibration and evaluation data are available in prior work [5] and 661k is already of sufficient size to be representative. We used the 305 genome batches that were inferred previously using MiniPhy [5], and embedded in the current version of Phylign (https://github.com/karel-brinda/phylign). To account for the effects of data quality and contamination, we considered both its complete version (661k, n=661,405), as well as its subset omitting genomes that failed quality control (661k-HQ, n=639,981). In particular, the impact of omitting low-quality genomes has previously been shown as major for computational trade-offs [5]. We benchmarked the indexes on a single cluster node, with 30 CPUs and 200 GB of memory available.

First, we focused on disk sizes (**Fig. 2a**). The figure shows a consistent pattern across both collections. COBS produces by far the largest subindexes, even after XZ compression: its compressed footprint remains roughly an order of magnitude above all other tools. Fulgor and Fulgor-m form the smallest group overall, with XZ compression yielding only marginal additional savings. Themisto-h and Themisto-r occupy an intermediate band, with moderate reduction under XZ compression. Metagraph’s BRWT variant is extremely small even when uncompressed, suggesting that most structural redundancy has already been removed by the BRWT transform. The absolute values further highlight the divergence: across all 305 batches, COBS without compression is in the multi-terabyte range, Themisto around a few hundred gigabytes, and Fulgor consistently below that. Interestingly, all indexes except Metagraph have a similar size when they are compressed.

**Fig. 2.**
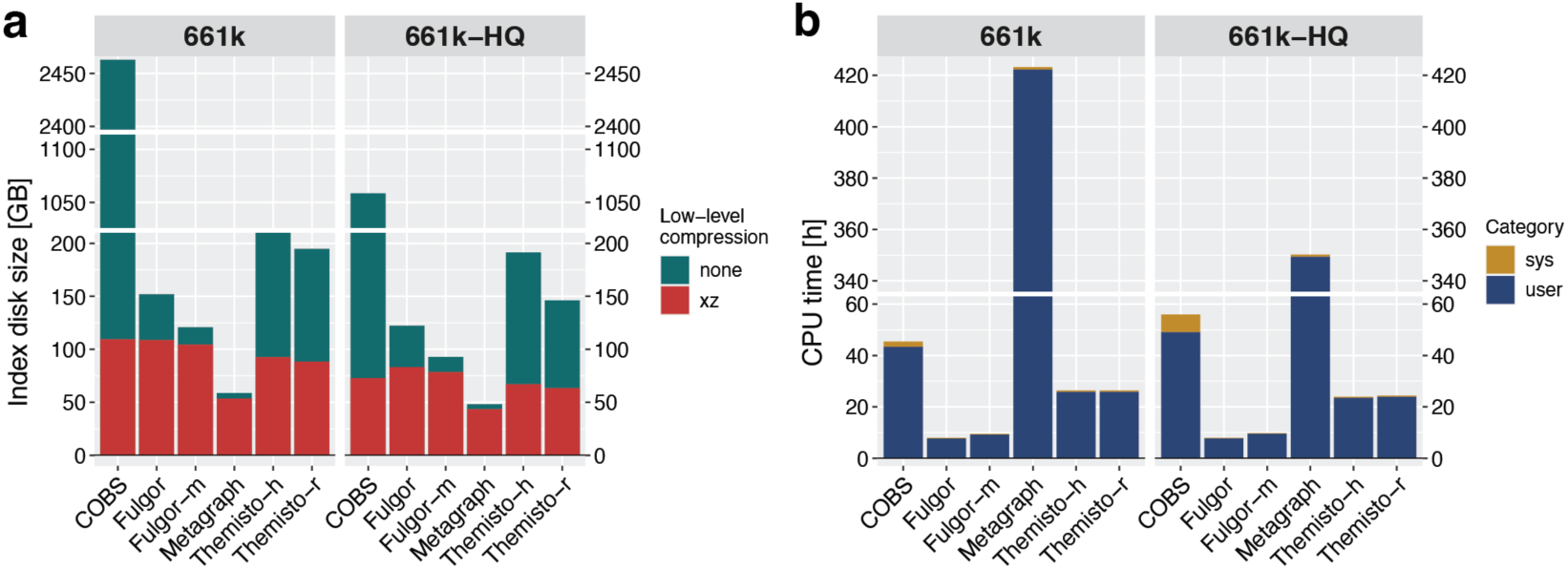
Cumulative disk and time requirements of COBS, Fulgor, Metagraph, and Themisto *k*-mer subindexes. Bars summarize results across all 305 phylogenetic batches in the 661k and 661k-HQ collections. **a) Disk sizes**. Total on-disk footprint of the *k*-mer subindexes, stored either uncompressed (green) or XZ-compressed (red), aggregated across 305 batches. **b) Search time for the 2,826 EBI plasmids.** Total CPU time (user + system) required for *k*-mer matching, summed across all batches. All thresholds were set to t=40% (**Supplement**).

Second, we sought to understand differences in query performance at the level of individual indexes (**Fig. 2b**). Here, we observed substantial differences, and clear separation according to index types. The clear winner was Fulgor, which – both base and meta-colored – dominates with the lowest total CPU time across both 661k and 661k-HQ. Themisto is consistently ∼2× slower than Fulgor, regardless of variant. COBS is another ∼2× slower than Themisto, confirming that its bit-sliced structure is penalized under heavy *k*-mer enumeration. Metagraph is an extreme outlier: even its optimized BRWT configuration is more than an order of magnitude slower than the others, reflecting the cost of decompression and traversal of a deeply compressed structure.

#### Batch-level heterogeneity in space–-time trade-offs

To better understand index behavior beyond global averages, we examined performance on the 305 phylogenetic batches of the 661k-HQ dataset (**Fig. 3**). A major advantage of the subindex paradigm is that contributions of individual subindexes can be perfectly isolated and studied in isolation. We reused the data from the previous experiment, but slightly readjusted the formulation. As all indexes except COBS were sufficiently compressed on their own, we proceeded with seven subindex configurations: Fulgor, Fulgor-m, Themisto-r, Themisto-h, Metagraph, COBS, and COBS-xz (**Supplement**). In the latter (COBS-xz), we assume that the index is stored on disk in XZ-compressed form and decompressed on the fly, as implemented in Phylign.

**Fig. 3.**
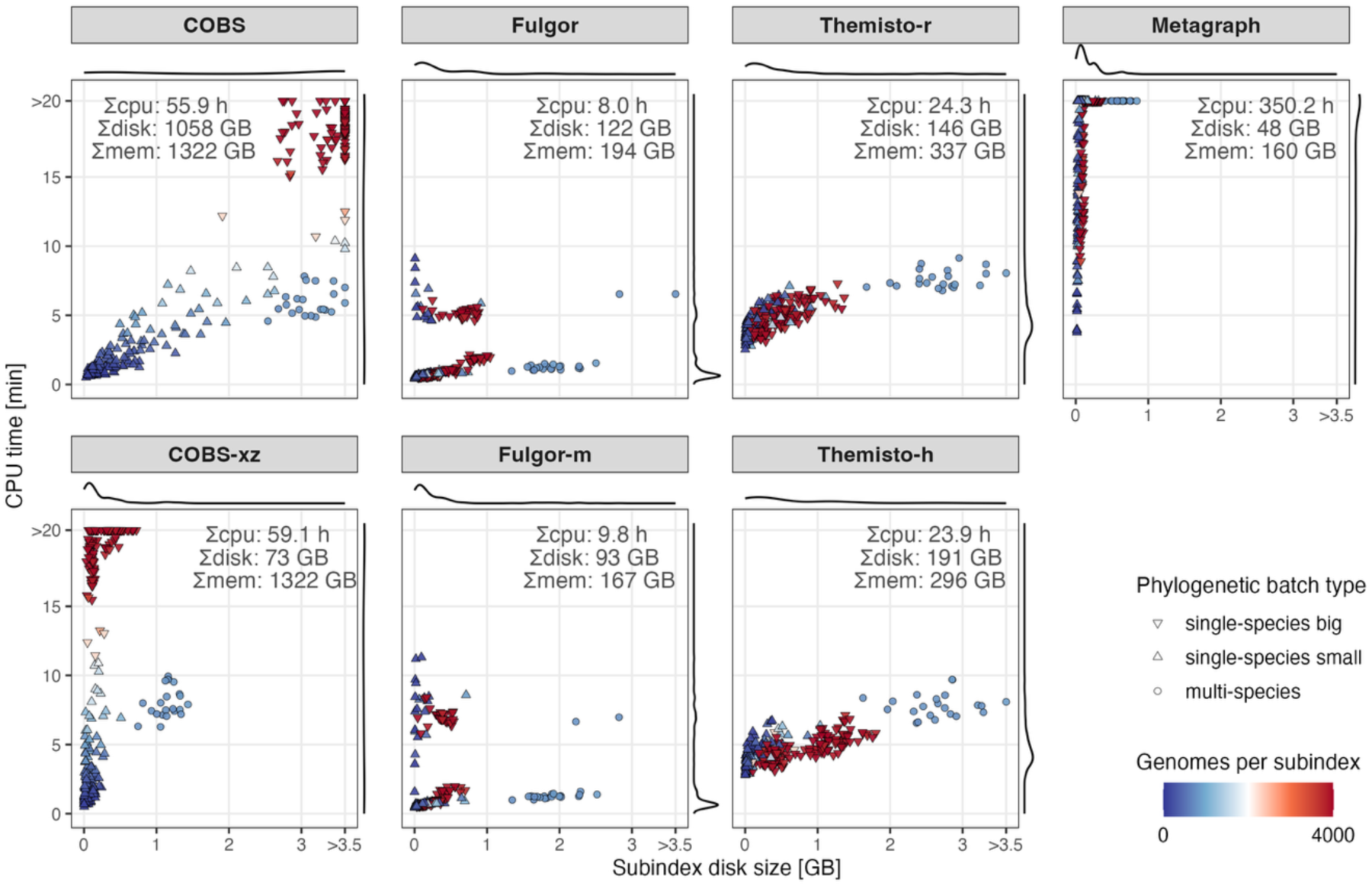
Batch-level performance of *k*-mer subindexes in the 661k-HQ plasmid search. For each index type, each point represents one phylogenetic batch; the x-axis shows the on-disk subindex size (GB), and the y-axis shows CPU time (min) to query the EBI plasmid set (same experiment as in Fig. 2). Point color encodes the number of genomes per subindex, and point shape distinguishes large single-species, small single-species, and multi-species (“dustbin”) batches (threshold: 2,000 genomes). Text in each panel reports the total CPU time, total disk footprint, and total memory footprint across all batches. Marginal densities summarize the distributions of CPU time and disk size.

The 661k-HQ collection contains 639,981 high-quality genomes from 2,336 species, with just 20 species accounting for 90% of genomes [3]. After phylogenetic compression, the top ten species – nearly 80% of the genomic content – occupy less than half of the database space, whereas “dustbin” batches of sparsely sampled species expand to proportions 9.4× larger than their precompression share [3]. To reflect this structure, MiniPhy defines 305 batches for 661k-HQ, of which 283 are single-species and 22 are multispecies dustbin batches [3]. In **Fig. 3**, we further split single-species batches into large (≥2,000 genomes) and small ones, and examined disk–time trade-offs separately for these three batch classes.

We found that the index performance is strongly batch-dependent (**Fig. 3**). For all indexes, we see batches of the same type clustered. Large, low-diversity single-species batches tend to be best served by Fulgor/Fulgor-m, which are both small and fast, whereas high-diversity dustbin and small-species batches often favor COBS-xz, which remains time-competitive at modest size thanks to phylogenetic compression. Metagraph provides by far the best compression, but at the price of uncompetitive query times. Consistent with recent *k*-mer matching literature [42], all tools use substantially more memory compared to disk index size. No single index dominates across all phylogenetic batches, which motivates mixed-index optimization.

#### Pareto optimization of mixed index configurations

The large differences across index types prompted our interest in whether combining multiple index types within the same workflow provides measurable benefit. For instance, **Fig. 3** suggests that high-diversity batches might benefit from COBS, whereas low-diversity ones could be served more efficiently with Fulgor. This can be formulated mathematically as follows: for a given disk space budget, how should we assign index types to individual batches so that they fit within the budget and simultaneously minimize the search time? This yields a standard multi-objective combinatorial optimization problem in the Pareto sense.

Using the batch-level disk–time measurements, we computed the global Pareto front (**Fig. 4a**). For each batch and index type, we considered a cost vector (disk size, CPU time). First, we created two-objective Pareto sets for each batch, removing batch-level dominated configurations. Then, we computed the exact global Pareto front over all assignments by iteratively combining batches: starting from the zero-cost point (0 disk, 0 time), we repeatedly took the Minkowski sum of the current front, with the local Pareto set of the next batch and pruned globally dominated points. This yields the non-dominated combinations of index types over all batches under additive disk and time costs, in the standard setting of multi-objective combinatorial optimization [43]. Conceptually, this procedure performs repeated Pareto sums of the batch-level Pareto sets, followed by dominance filtering, as studied for unions and Minkowski sums of non-dominated sets [44]. From these data, we also extracted three extreme assignments (**Fig. 4b**): the global minimum-time solution (fastest index for each batch), the global minimum-size solution (smallest index for each batch), and the minimum-size solution omitting Metagraph.

**Fig. 4.**
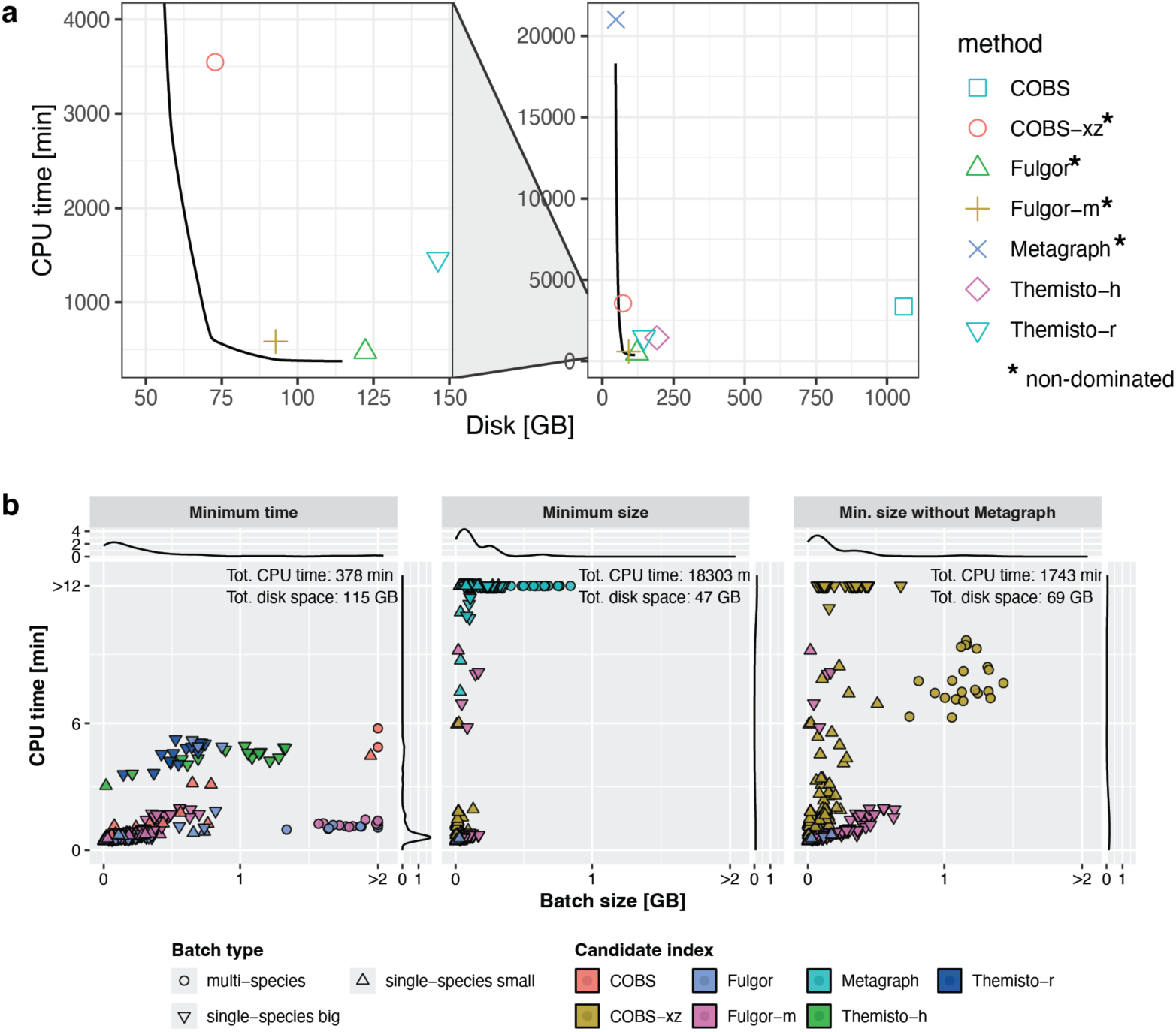
Pareto optimization of plasmid subindex search in 661k-HQ. **a) Pareto optimization analysis.** Global Pareto front for joint index-type assignment, plotted together with points corresponding to single-type indexes. **b) Extreme assignments.** Index types and performance characteristics of subindexes at the three Pareto assignments (minimum-time, minimum-size, and minimum-size-without-Metagraph).

The examination of the front shows a characteristic steep descent: aggressive space minimization rapidly incurs extreme time penalties. The practical L-shaped region of the curve is dominated by three closely clustered candidates – COBS-xz, Fulgor-xz, and Fulgor – all lying near the knee of the efficient front and offering the strongest real-world trade-offs.

We then examined the index composition of the Pareto front endpoints (**Fig. 4b**). The fastest achievable solution required approximately 6 h and consumed 115 GB of disk space (**Fig. 4b**, left panel). Conversely, minimizing disk space reduces the requirements to 47 GB, but at a prohibitive cost of approximately 304 h of search time (**Fig. 4b**, central panel). Because these extremely slow configurations were consistently driven by Metagraph, we excluded all solutions containing Metagraph from further consideration.

Recomputing the Pareto front without Metagraph produced a more practical landscape. Disk requirements for indexing can be reduced to 69 GB of disk space (**Fig. 4b**, right panel), requiring a total of approximately 26 h of search time, while the minimum search time remained unchanged relative to **Fig. 4b** (left panel). Among all single-type indexes, only COBS-xz and Fulgor-m fall between the two Pareto endpoints, making them the strongest compromises between size and speed in practice: COBS is consistently smaller, whereas Fulgor is consistently faster.

### Phylign-Fulgor and its experimental evaluation

#### Implementation and calibration

Once subindex benchmarking and optimization are complete, the final step is to integrate the indexes into the target workflow such as Phylign. This can be done either via a single index type, which is technically easier but less efficient, or as a combination of multiple index types. As Phylign does not currently support a simultaneous use of multiple index types, we focused on a single-index configuration. Based on our analyses, we selected Fulgor-m (meta-colored Fulgor) as the base index. Due to Fulgor’s limited native support for subindex search (**Tab. 1**), we created its custom modification (a fork of v2.0.0), adapted to the subindex paradigm, with modified parameters, a more suitable output format, and an improved control over *k*-mer matching (**Supplement)**.

We developed Phylign-Fulgor, a fork of the original Phylign embedding Fulgor. This required a series of source code modifications (**Supplement**). We included both the 661k-HQ (n=639,981 genomes, 305 batches) and ATB-HQ collections (release v0.2, n=1,858,610 genomes, 650 batches). In both cases, phylogenetically compressed Fulgor-m indexes were generated analogously to the previous benchmarks (Supplement). Phylign-Fulgor is provided under the MIT license at https://github.com/Francii-B/Phylign-Fulgor, and all database files are available on Zenodo (**Tab. S1**)

Phylign-Fulgor provides substantial dynamic range in its search sensitivity, primarily controlled by the *k-*mer matching threshold parameter. To make its results comparable with LexicMap and Phylign-COBS as the closest methods, we identified those thresholds that equalized for a single plasmid the number matched genomes (**Fig. 5**). First, we used ATB-HQ to identify the maximum number of detectable matches by Phylign-Fulgor, which was 498,410 corresponding to all sequences sharing at least one *k*-mer with the query (t≈0.00005%). Then, we used ATB-HQ collection to identify a threshold matching LexicMap’s count (485,281 matches): this was achieved via setting a threshold t=0.007%, yielding 481,499 genomes (**Fig. 5**). Finally, we performed an analogous calibration on the 661k-HQ collection to reproduce the number of genomes matched by Phylign-COBS (with its previously recommended threshold t=40%): this yielded 2,949 matches, back-translated to Phylign-Fulgor threshold 18% with 2,982 matches (**Fig. S1**).

**Fig. 5.**
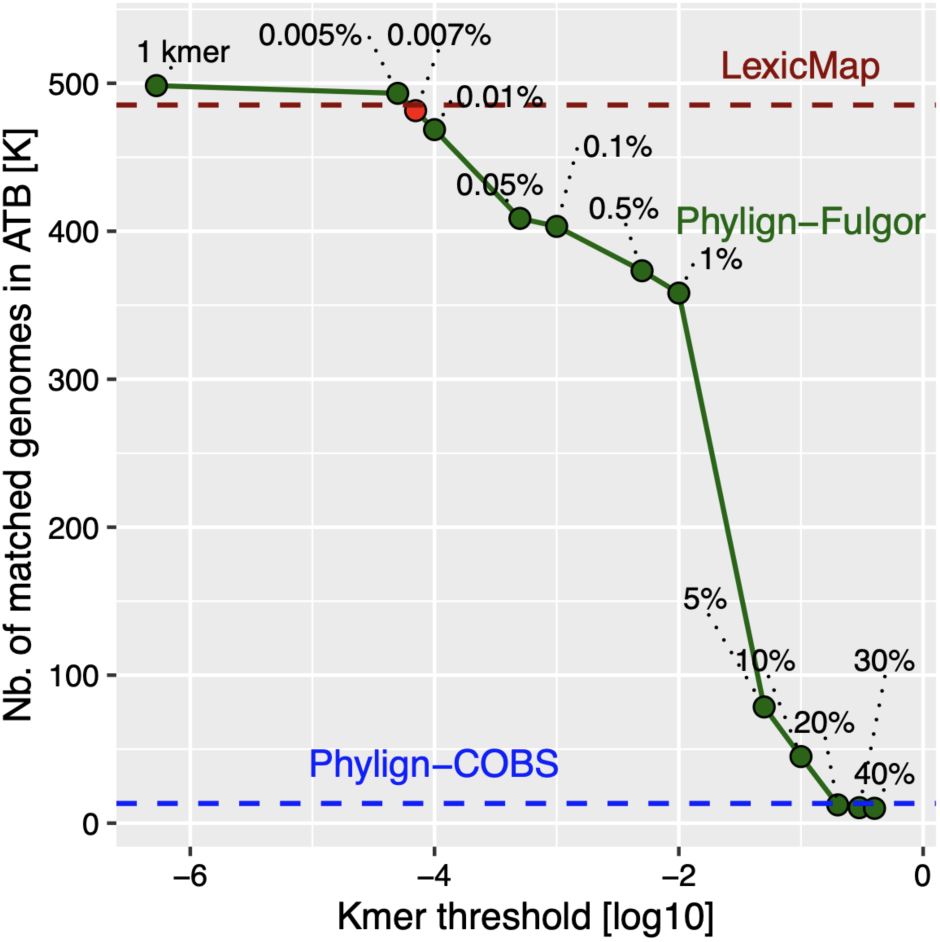
Dynamic range of Phylign-Fulgor. The graph shows the number of matched genomes for the pCUVET18-1784 plasmid in the ATB-HQ collection as a function of the minimum required proportion of matching *k*-mers in the *all-matches-above-threshold* mode, compared with LexicMap and Phylign-COBS (threshold t=40%, as lower values cause a combinatorial explosion in COBS). The leftmost point corresponds to requiring a single matching *k*-mer; the red point (t=0.007%) matches the LexicMap-calibrated threshold. The blue line reports Phylign-Fulgor results using across the ATB-HQ collection using the threshold calibrated relative to Phylign-COBS (t=18%, n=13,302).

#### Experimental design

We compared Phylign-Fulgor to the two existing state-of-the-art tools locally capable of alignment to million-genome collections: Phylign [5] and LexicMap [22] (**Figs. 6, 7**). To evaluate the impact of replacing COBS with Fulgor within Phylign, we assessed performance under two comparable experimental settings. First, we reproduced on a standard laptop the plasmid-search experiment described in the Phylign paper [5], querying the EBI plasmids against the 661k-HQ collection. Second, we queried the pCUVET18-1784 plasmid across the same 661k-HQ dataset to compare the sensitivity of Phylign and Phylign-Fulgor. Similarly, we evaluated the performance of Phylign-Fulgor and LexicMap by replicating the plasmid-search experiment from the LexicMap paper [22], querying the pCUVET18-1784 plasmid against the ATB-HQ collection, and we additionally queried first 100 plasmid from the same EBI dataset to assess performance across both laptop and server environments..

**Fig. 6.**
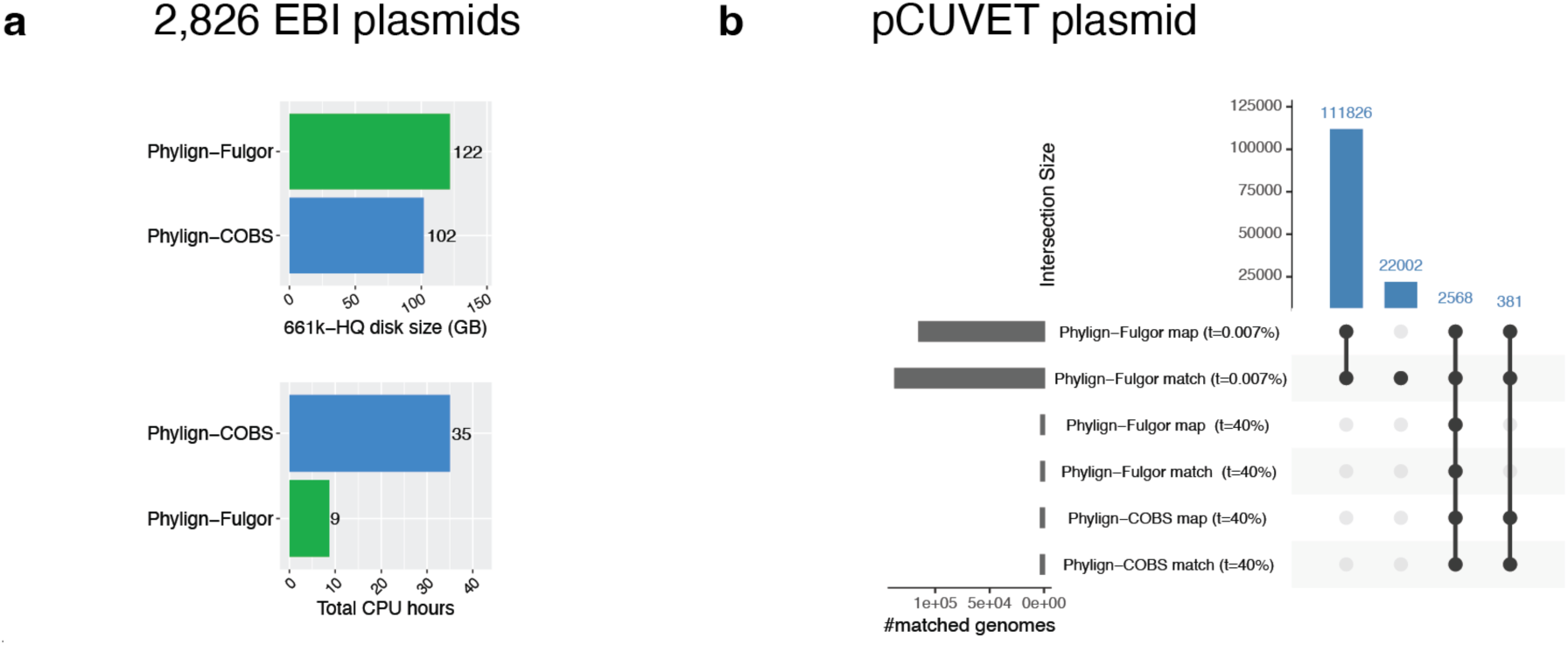
Comparative performance of Phylign-Fulgor and Phylign-COBS on the 661k-HQ collection. **a)** Database on-disk size and CPU time for k-mer matching of the 2,826 EBI plasmids on a laptop. The following parameters were set for both Phylign-COBS and Phylign-Fulgor: *kmer_thres: 0.4; nb_best_hits: 1000; minimap_preset: asm20.* Search and matching statistics are reported in **Tab. S2**. **b)** UpSet plot showing intersections of matched genomes between methods for the pCUVET18-1784 query. Phylign-Fulgor was run with two thresholds (t=0.007% and t=40%) and Phylign-COBS with t=40%. Both programs were run in the following setup: *nb_best_hits: 2000000; minimap_preset: asm20*.

#### Comparative performance of Phylign-Fulgor and Phylign-COBS

First, we evaluated how the *k*-mer matching step in Phylign is affected by replacing COBS with Fulgor (**Fig. 6a, Tab. S2**). We used the *k*-mer matching experiments from the BIGSI benchmark [17], which was also the first step of the original Phylign benchmark [5], and queried the 2,826 EBI plasmids against the 661k-HQ collection on a laptop.

Phylign-Fulgor performed *k*-mer matching approximately 4× faster than Phylign-COBS (8.4 h vs. 35 h of total CPU time), with a 1.2× increase in its index size (from 102 GB to 122 GB) (**Fig. 6a**). However, because Fulgor implements exact *k*-mer indexing, it detected a factor of 2.5× fewer matches than COBS (3,732,698 vs. 9,163,123) (**Tab. S2**). This reduction did not substantially affect the runtime of the subsequent alignment step (Phylign-Fulgor: 14 CPU hours; Phylign-COBS: 15 CPU hours), but it did decrease the total number of alignments (7,705,870 vs. 8,980,408).

We next assessed how the COBS-to-Fulgor replacement affects sensitivity. For this analysis, we queried the pCUVET18-1784 plasmid across the 661k-HQ collection, reporting all detected matches (*nb_best_hits: 2000000*), and compared the sets of matched genomes obtained before threshold calibration (t=40%) and after calibration with respect to LexicMap (t=0.007%).

Using the threshold previously recommended for Phylign-COBS (t=40%), both tools identified a comparable number of matched genomes (2,568 for Phylign-Fulgor vs. 2,949 for Phylign-COBS), with all Phylign-Fulgor matches also found by Phylign-COBS (**Fig. 6b**). When the calibrated high-sensitivity threshold was used for Phylign-Fulgor (t=0.007%), it detected 136,777 matches (approximately 46× more than Phylign-COBS at t=40%) while still recovering all matches detected by Phylign-COBS.

Despite the large variation in match counts across thresholds, all Phylign-Fulgor *k*-mer matching runs completed in 1–2 minutes of real time (4–6 minutes of total CPU time), making them 4–8× faster than Phylign-COBS in real time and 30–48× faster in total CPU time (Phylign-COBS real time: 10 minutes; total CPU time: ∼2 h) (**Tab. S3**).

As the number of *k*-mer matches directly determines the workload of the subsequent alignment step, we also assessed how the increased match count affects alignment performance. With thresholds t=40%, Phylign-Fulgor was ∼2× faster than Phylign-COBS (∼10 min vs. 18 min of real time; ∼1.5 h vs 3.5 h of total CPU time), with only a modest reduction in detected alignments (22,956 vs 25,870) (**Tab. S3**).

#### Comparison to LexicMap on AllTheBacteria

As LexicMap is the current recommended tool for searching in AllTheBacteria, we sought to compare Phylign-Fulgor and LexicMap performance in a comparable setup. We evaluated two query types across the ATB-HQ collection: (1) the pCUVET18-1784 plasmid, replicating the plasmid-search experiment from the LexicMap paper [22], and (2) a batch of the first 100 plasmids from the same dataset. Because LexicMap is not scalable to portable devices, all LexicMap experiments were performed on a server node with an SSD disk and 20 CPUs available, whereas Phylign-Fulgor experiments were performed both on the server and additionally on a standard laptop, using its threshold calibrated with respect to LexicMap (t=0.007%).

In contrast to the original Phylign (Phylign-COBS), where *k*-mer matching and alignment required similar amounts of time [5], Phylign-Fulgor’s *k*-mer matching step was hundreds of times faster than the subsequent Minimap2 alignment step (∼830× for the single plasmid and ∼180× for the 100-plasmid batch). This is thanks to Fulgor’s high throughput and sensitivity, which generate many more matches in much less time.

When compared to LexicMap, Phylign-Fulgor *k*-mer matching was ∼3.1× faster for the single plasmid and ∼28× faster for the 100-plasmid batch, whereas the full Phylign-Fulgor workflow (*k*-mer matching + Minimap2 alignment with the “asm20” preset) was ∼14× slower for the single plasmid but 1.4× faster for the 100-plasmid batch (**Fig. 7a**). This reflects the design of Phylign: it iterates over all phylogenetically compressed assemblies (unlike LexicMap, which keeps all assemblies uncompressed) and therefore has higher single-query costs, but these amortize once the batch is larger.

**Fig. 7.**
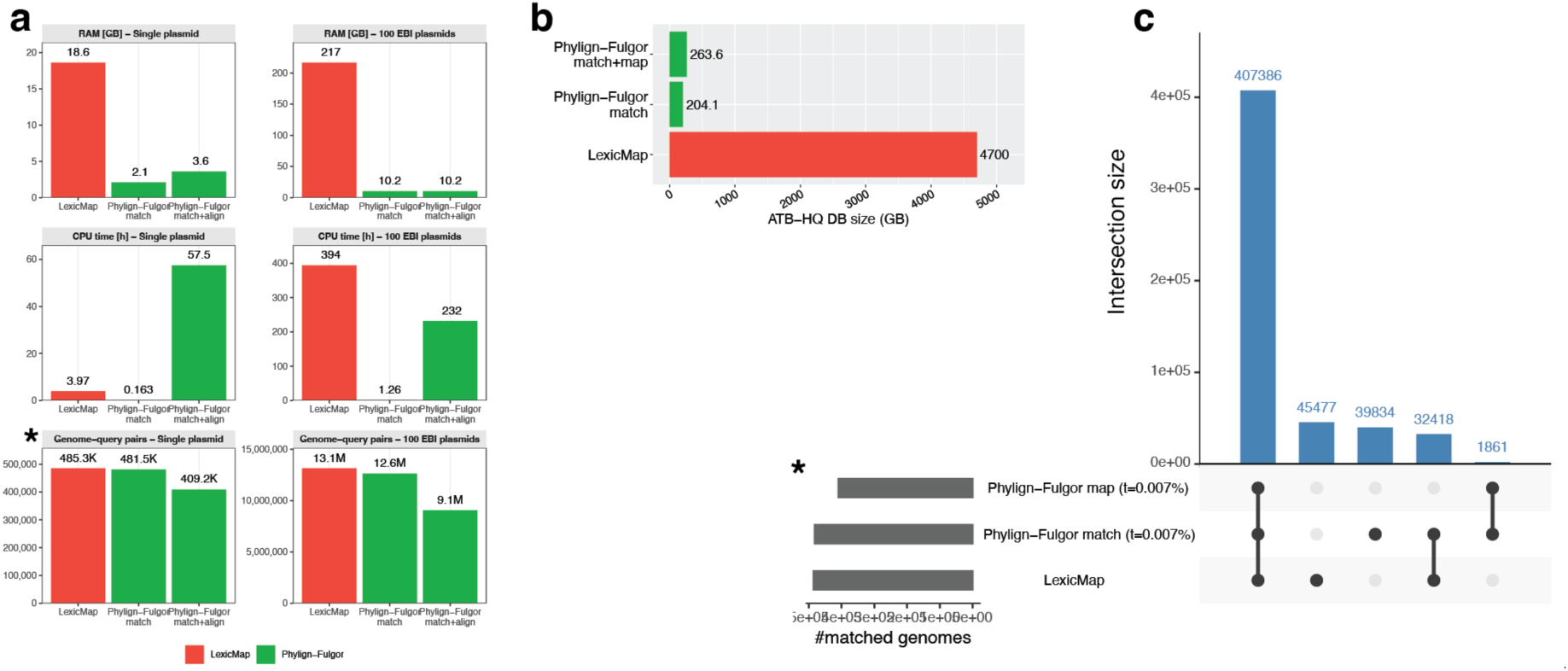
Comparative performance of Phylign-Fulgor (t=0.007%) vs LexicMap on the ATB-HQ collection. **a)** CPU time, RAM usage, and genome–query counts when searching the pCUVET18-1784 plasmid as a single query or a batch of 100 EBI plasmids. Phylign–Fulgor (match and match+map modes) requires substantially less RAM and disk space than LexicMap while returning comparable query counts. **b)** Total index size of each method on the ATB-HQ dataset. Phylign–Fulgor produces a ∼20× smaller index than LexicMap. **c)** UpSet plot showing intersections of matched genomes between methods for the pCUVET18-1784 query. The largest intersection corresponds to genomes detected by all methods, confirming concordance between Phylign–Fulgor and LexicMap. The benchmark was performed on a computing cluster using a server (20 CPUs, SSD disk, and 150 GB of available RAM) and a laptop (MacBook Pro 18,3 Apple M1 Pro, 10 CPU cores, 16 GB RAM). Phylign-Fulgor parameters: *nb_best_hits: 2000000; minimap_preset: asm20.* Detailed information is provided in **Tab. 2**.

**Table 2.**
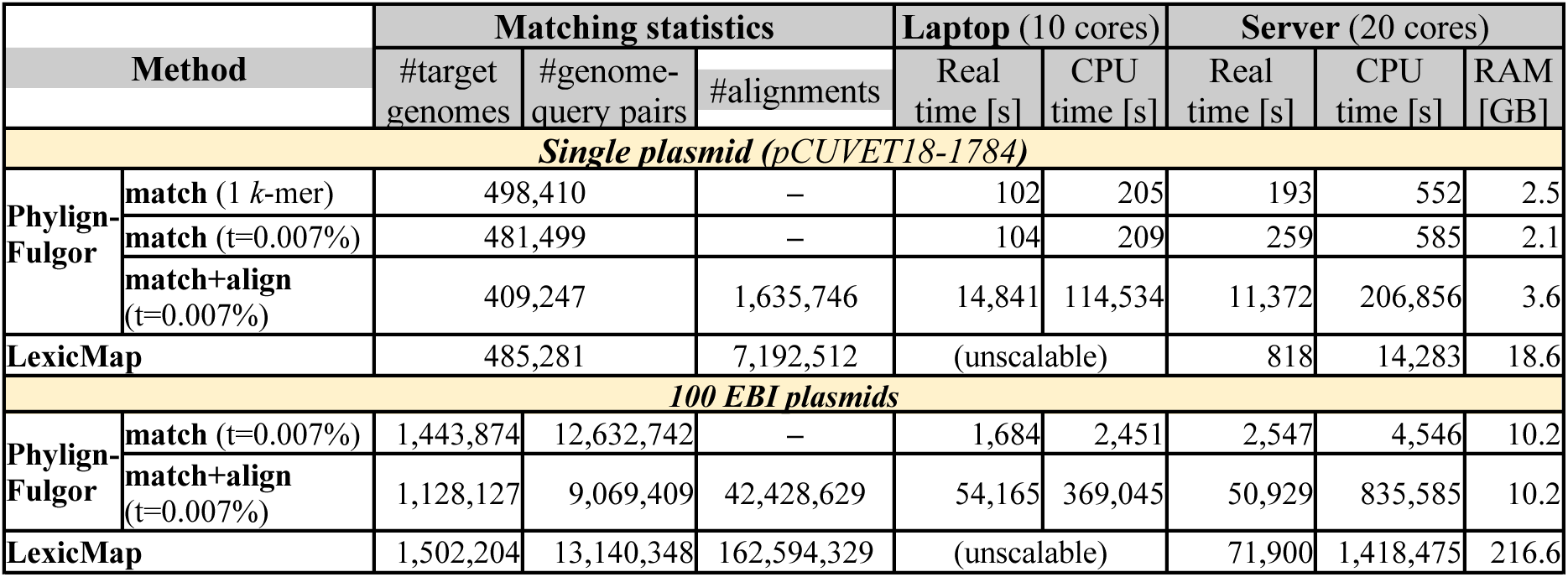
Detailed statistics for plasmid search in the ATB-HQ collection. The configuration of the computers and search parameters are provided in the legend of **Fig. 7**.

We then focused on Phylign-Fulgor space requirements in both memory and disk. On disk, its *k*-mer index required 204 GB, and the corresponding genome assemblies required ∼59 GB, making its overall space requirements 17–23× smaller than those of LexicMap (index size: 4.7 TB) (**Fig. 7b**). Phylign-Fulgor searches required substantially less memory than LexicMap, with reductions of 5–8× for single-plasmid searches and 21× for the 100-plasmid batch (**Fig. 7a**).

Finally, we compared the sensitivity of both methods (**Fig. 7a,c**; **Tab. 2**). Phylign-Fulgor *k*-mer matching detected a comparable number of query–genome pairs to LexicMap for both the single plasmid (481,499 vs. 485,281) and 100 plasmids (12,632,742 vs. 13,140,348) (**Fig. 7a**). After Minimap2 alignment, the number of query–genome pairs detected by Phylign-Fulgor decreased by 15–30%, resulting in ∼1.3× fewer pairs than LexicMap for both the single plasmid (n=409,247) and the 100-plasmid batch (n=9,069,409). To assess sensitivity differences, we compared the set of genomes detected by Phylign-Fulgor and LexicMap for the single plasmid. Among the 481,499 matched genomes, Phylign-Fulgor identified 90% (n=439,804) of those detected by LexicMap (**Fig. 7c**).

Overall, Phylign-Fulgor trades a small loss in alignment-level sensitivity for dramatic improvements in space usage (20× smaller), memory (5–21× lower), and *k*-mer filtering throughput (3–28× faster), while becoming competitive or faster than LexicMap for realistic batches.

## DISCUSSION AND CONCLUSIONS

Modern collections of millions of bacterial genomes may revolutionize point-of-care applications, yet the lack of local methods for alignment using portable devices is a critical gap. Here, we focused on *k*-mer-based search and developed a practical end-to-end methodology for combining modern *k*-mer indexes with phylogenetic compression [5] in local search via subindexes. We plugged this into Phylign, obtaining the Phylign-Fulgor pipeline, and showed that it can search such large collections on laptops within several minutes. As *k*-mer indexes have so far been mostly studied in isolation and applied on a large scale only on servers, this represents a major methodological advance.

One key observation is that index selection is not a single-metric choice, but a Pareto optimization problem balancing space, latency, batch behavior, and compatibility with the subindex paradigm. While this is intuitive, to the best of our knowledge, this is the first time to show this quantitatively. Across our plasmid-search use case, Fulgor, COBS, Themisto and Metagraph exhibited surprisingly different performance at the batch level depending on phylogenetic relatedness and batch size, reinforcing that cumulative high-level numbers for entire collections are not informative of index differences. Furthermore, most indexes are currently technically unsuited for the type of matching that would be needed, and source code modifications would almost always be necessary should they be used in practice.

Our study has several limitations. First, we relied on predefined MiniPhy batches, which fixes the phylogenetic granularity, which may not be optimal across all types of searches, taxonomic groups, or indexes. Given current compressive capabilities of Fulgor, one might imagine, for instance, building species-level indexes instead. Second, we used only the default parameter choice of each index, such as Fulgor’s minimizer size, yet these parameters may have a major effect on the resulting performance and could be easily included in the Pareto optimization. Third, we tested single type of queries in a constrained use case; yet other biological applications (e.g., metagenomic screening) may stress indexes in different ways. Finally, we did not evaluate the most recent Fulgor version, which substantially improved performance further, as this would require redoing our modification in its source code.

Future work will therefore focus on optimizing internal index parameters, co-optimizing batch construction with phylogenetic structure, and characterizing how species composition shapes size-speed trade-offs. We anticipate that the broader community may provide standard APIs in *k*-mer indexes, which would better allow us to integrate high-performance indexes in workflows such as ours without source code modifications. Ultimately, our goal is a fully adaptive version of Phylign with support for databases, which could select, per task, the optimal combination of indexes and parameters to achieve near-optimal search, with the goal of providing near-real time search across millions of genomes on commodity hardware. Given the accelerating growth of bacterial genome databases and the demand for local data analyses, the methodological principles established here – pragmatic index evaluation and pareto-guided selection, combined with phylogenetic compression – provide a foundation for the next generation of large-scale *k*-mer search tools.

## ACKNOWLEDGEMENTS

This research was supported by the French National Research Agency (ANR) under Grant ANR-24-CE45-1226 for the REALL project. Portions of this research were conducted at the GenOuest bioinformatics core facility (https://www.genouest.org).

## Supplement

### METHODS

#### Data and program preparation

##### 661k collection

All genomes from the 661k collection (n=661,309) [3] were downloaded from Zenodo using Phylign (command “*make download_asms*”) [5], as 305 phylogenetically compressed .tar.xz archives containing FASTA files. The corresponding Zenodo records are listed in **Tab. S1.** Phylogenetically ordered lists of genomes were generated for each batch and stored as text files using the following command:

> *tar -tf {batch}.tar.xz > {batch_related_genome_list}.txt*

HQ genomes were extracted from the 661k collection [3], based on its original metadata and analyzed as a separate collection. The HQ information was used as reported in *File4_QC_characterisation_661K.txt* (**Tab. S1**). The batches and phylogenetic orderings were used as defined in [5].

#### AllTheBacteria

Genomes of the ATB collection from release 0.2 (n=2,440,377) [4] were downloaded as 665 phylogenetically compressed .tar.xz archives of FASTA files from OSF (**Tab. S1**), together with their metadata stored in *file_list.all.20240805.tsv.gz.* Phylogenetically ordered lists of all genomes, as well as for the HQ subsets (n=1,858,610), were generated for each batch, following the same procedure described for the 661k collection. The HQ information was obtained from the file *hq_set.sample_list.txt.gz*, available on OSF.

#### Used indexes

We used four *k*-mer indexes selected for evaluations:

● Fulgor v. 2.0.0 [15, 25], the default and the meta-colored variants (Fulgor and Fulgor-m, respectively).
● Themisto v.3.2.2 [16], both hybrid and roaring bitmap variants (Themisto-h and Themisto-r, respectively).
● Metagraph v.0.3.6 [7] (row-diff relaxed Multi-BRWT variant).
● COBS v.0.3.1 [23].

All indexes were downloaded from their respective GitHub repositories (**Tab. S1**).

### Evaluation of index space requirements

Evaluations of index space requirements were performed on the GenOuest cluster (https://www.genouest.org), using a single node with 30 CPUs and 200 GB of memory available for computation.

For each considered index (except COBS, which was already evaluated in [5]), we developed a dedicated Snakemake [9] workflow to generate batch-level compressed indexes for each batch of the 661k and 661k-HQ collections (305 batches in both cases). All indexes were created such that the phylogenetic reordering is used at its max: the batches were phylogenetic and we retained the phylogenetic order in the input for each program.

The specific commands used for generating each subindex are reported below.

#### Fulgor and Fulgor-m

For each batch, we built both Fulgor index types [15, 25] via a single invocation of the following command:

> **fulgor build -k 31 -m 20 -l {batch_genome_list} -o {fulgor_index_name} --check --meta**

#### Themisto-h and Themisto-r

We built the Themisto [16] indexes using the following commands:

> **themisto build -k 31 -d 20 -i {batch_genome_list} -o {themisto_index_name} -s {annotation_type}**

where {*annotation_type}* indicated the specific type of data structure to use for the color annotation, namely “*roaring*” to build Themisto-roaring bitmap indexes (Themisto-r) or “*sdsl-hybrid*” to generate Themisto-hybrid indexes (Themisto-h).

#### Metagraph

We considered the Row-diff relaxed Multi-BRWT variant of Metagraph [7]. The following commands were used to generate the dBG, its annotation matrix and to convert this last into row-diff relaxed Multi-BRWT format:

> **cat {batch_genome_list} | metagraph build -k 31 --graph succinct --state small -o**

> **{metagraph_index_name}.dbg**

where *{metagraph_index_name}.dbg* corresponds to the output dBG.

> **cat {batch_genome_list} | metagraph annotate -i {metagraph_index_name}.dbg --anno-filename -o**

> **{metagraph_column_annotation}**

where *{metagraph_index_annotation}* corresponds to the file name of the output annotation matrix.

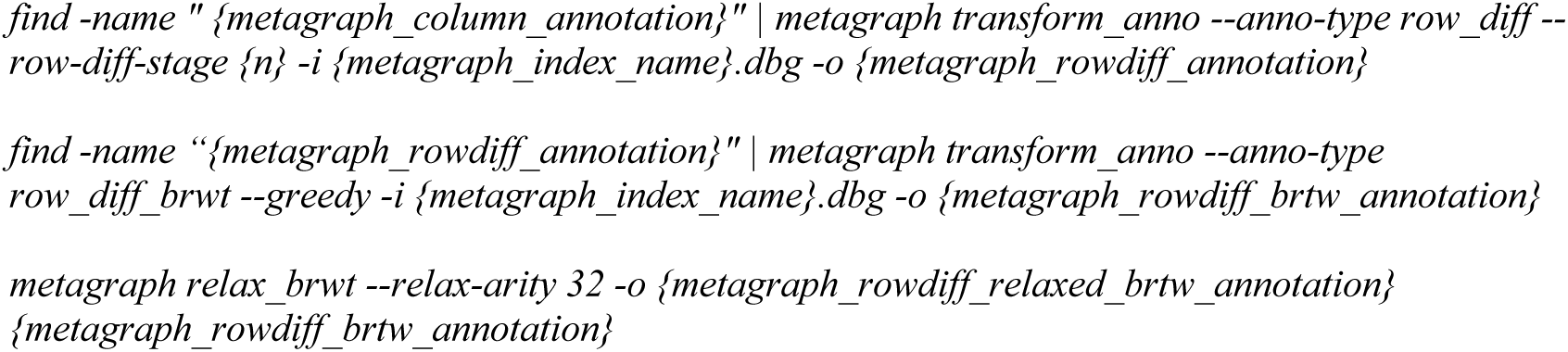

where *{metagraph_rowdiff_relaxed_brtw_annotation}* represents the final annotation file in the row-diff relaxed Multi-BRWT format.

The generated subindexes were subsequently compressed using the the low-level compressor XZ, using the following command:

> **xz -9 -T1 -k {uncompressed_subindex}**

In order to calculate the overall collection size for each index, before and after compression, individual subindex sizes were calculated as follows:

> **stat -c "%s %n" {XZ-compressed/uncompressed_subindex}**

### COBS

Phylogenetically compressed COBS indexes of the 661k collection and its HQ subset are publicly available on Zenodo (**Tab. S1**). Each COBS subindex of the 661k-HQ collection was downloaded from the corresponding Zenodo records using Phylign (command “*make download_cobs*”), while COBS subindexes of the entire 661k collection were downloaded modifying the specific https address of the Zenodo record in Phylign code and then downloaded using the same command.

### Evaluation of search time

Evaluations of search time were performed on the GenOuest cluster, using a single node with 30 CPUs and 200 GB of memory available for computation. Searches were performed using the EBI plasmid collection (n=2,826) (**Tab. S1**) as queries. Plasmid sequences were downloaded using their corresponding plasmid IDs as follows:

> **curl -o {plasmidID}.fasta "*https://www.ebi.ac.uk/ena/browser/api/fasta/{plasmidID}"*

and stored in a single Multi-FASTA file.

For each index type, we created a dedicated Snakemake workflow to perform *k*-mer matching of the EBI plasmids across the 661k collection and its HQ subset, searching for all matches carrying at least 40% of query *k*-mers via commands below. XZ-compressed indexes were first decompressed as follows:

> **xz --decompress -k {XZ_compressed_index}**

Uncompressed indexes were queried individually. Decompression and search time was measured separately for each batch via GNU Time via Galitime (**Tab. S1**). The specific commands used for *k*-mer matching are reported below.

### COBS

> **cobs query -i {cobs_subindex} -f {EBI_plasmids} -t 0.4 -T 4 --load-complete > {output_matches}**

#### Fulgor and Fulgor-m

> **fulgor pseudoalign -i {fulgor/meta-fulgor_subindex} -q {EBI_plasmids} -o {output_matches} --threshold 0.4 -t 4**

where *{meta-fulgor/fulgor_subindex}* could be either a fulgor or a meta-fulgor index.

#### Themisto-h and Themisto-r

> **themisto pseudoalign -i {themisto_subindex} -q {EBI_plasmids} -o {output_matches} --threshold 0.4 --include-unknown-kmers -t 4**

where *{themisto_subindex}* could be either a Themisto-hybrid or a Themisto-roaring bitmap index, while “*--include-unknown-kmers”* specifies that similarities should be computed considering all query *k*-mers.

#### Metagraph

> **metagraph query -i {metagraph_subindex}.dbg -a {metagraph_subindex}.annodbg --count-labels --discovery-fraction 0.4 -p 4 --fwd-and-reverse {EBI_plasmids}**

where *{metagraph_subindex}.dbg* and *{metagraph_subindex}.annodbg* represented the graph and annotation of each Metagraph subindex, respectively.

### Phylign-Fulgor implementation and comparisons

#### Modified Fulgor

To enable compatibility with the subindex paradigm and to integrate it in the Phylign workflow, we modified the Fulgor source code (release 2.0.0, commit 12da1e9). This modified version was called “modified-Fulgor” (https://github.com/Francii-B/modified-Fulgor)(Tab. S1).

● **Adjusting *k-*mer matching to account for all query *k*-mers.** Fulgor implements the so-called *threshold-union* regime of *k*-mer matching to identify *all-matches-above-threshold*, where matches are returned if they share a minimum proportion of *k*-mers with the query. For each match, this proportion is computed by considering only positive *k*-mers, by default. Therefore, Fulgor’s code was modified to compute this proportion considering the total number of query *k*-mers (modified-Fulgor, commit 7b040c4).
● **Returning the number of shared *k*-mers in the output.** Fulgor’s code was modified in order to return the output in a TSV format, providing query and match IDs, along with their corresponding number of shared *k*-mers (modified-Fulgor commit 7b040c4). Matches are output in descending order based on their number of shared *k*-mers (modified-Fulgor commit cd2c2fa).
● **Formatting output in a COBS-like format.** To integrate Fulgor into the Phylign workflow and enable the identification of best-matching genomes across different batches, the output file structure was modified to follow a COBS-like format (modified-Fulgor, commit 4a6c366). To allow users to select this output format, the “--cobs” flag was added among the input parameters (modified-Fulgor, commit 7b040c4).
● **Returning all matches sharing at least one *k*-mer.** Modifications were introduced to Fulgor’s code to set to 1 the minimum number of *k*-mers required to report matches when the similarity threshold is set to 0 (modified-Fulgor, commit 669536e).

#### Phylign-Fulgor

To replace COBS with Fulgor, the Phylign repository (https://github.com/karel-brinda/phylign, commit e1a2115) was forked to create Phylign-Fulgor (https://github.com/Francii-B/Phylign-Fulgor). Modified-Fulgor v. 2.1.0 was integrated into Phylign as an external submodule (Phylign-Fulgor, commit 04962db).

Phylign-Fulgor searches were implemented for both the 661k collections (main branch) and for the ATB collection (*ATB* branch). Meta-Fulgor indexes for the ATB collection were generated following the same procedure as described for the 661k collection. Integration of Fulgor required the following changes in Phylign:

● **Reference collection download command.** The Fulgor-m uncompressed indexes of the 661k and ATB collections were uploaded to Zenodo (**Tab. S1**). To enable downloading of Fulgor-m batches of the 661k collection, download links were replaced with those of the Fulgor-m indexes both in the main (Phylign-Fulgor, commit a3f915e) and in the *ATB* branch (Phylign-Fulgor, commit 7dbb010).
● **Removal of index decompression step.** The index decompression step was eliminated as uncompressed Fulgor-m indexes were integrated in Phylign (Phylign-Fulgor, commit df2d7c4).
● **Perform *k*-mer matching using Fulgor**. COBS *k*-mer matching command was replaced with Fulgor’s corresponding command, adjusting prefixes and suffixes in filenames for referring to Fulgor indexes (Phylign-Fulgor, commit df2d7c4).

#### Sensitivity calibration

To calibrate sensitivity of Phylign-Fulgor with respect to Lexicmap and Phylign-COBS, *k*-mer matching was performed to identify the thresholds that produced comparable numbers of matched genomes when searching the pCUVET18-1784 plasmid from *Serratia nevei* strain CUVET18-1784 (GenBank accession: CP115019.1; length: 52,830 bp). Specifically, Phylign-Fulgor was calibrated with respect to the number of genomes matched by Phylign-COBS across the 661k-HQ collection, benchmarking thresholds ranging from 5% to 40%, and with respect to the genomes aligned by LexicMap across the ATB-HQ collection, benchmarking thresholds ranging from 0% to 40%.

#### Comparison with the original Phylign (Phylign-COBS)

To evaluate the impact of replacing COBS with Fulgor in the Phylign, both Phylign-Fulgor and Phylign-COBS were benchmarked on a MacBook Pro 18,3 (Apple M1 Pro, 10 CPU cores, 16 GB RAM). Searches were performed using the EBI plasmid collection against both the 661k-HQ collection, using the following parameters: *kmer_threshold*=0.4, *nb_best_hits*=1000, *minimap_preset*=asm20.

Sensitivity assessments were performed by searching the pCUVET18-1784 plasmid across the 661k-HQ collection. Specifically, Phylign-COBS was benchmarked setting a threshold of 40% while Phylign-Fulgor was tested using both its uncalibrated (40%) and calibrated thresholds (0.007% and 18%). For both Phylign-Fulgor and Phylign-COBS, the following parameters were set: *nb_best_hits:* 2000000*, minimap_preset:* asm20*, threads:* 20*, max_ram_gb:* 80. These experiments were executed on the GenOuest cluster, using a single node with an SSD disk, 20 CPUs, and 80 GB of RAM available.

#### Comparisons with LexicMap

LexicMap index [10] for the ATB-HQ collection was built on the GenOuest cluster using the following command:

> **lexicmap index -S -X {HQ_ATB_genome_list} -O {lexicmap_index} -b 15000 --log {log_file}**

Searches with LexicMap and Phylign-Fulgor were performed both on the GenOuest cluster (single node with an SSD disk, 20 CPUs, and 300 GB of RAM) and on the laptop described in the previous section. The pCUVET18-1784 plasmid, previously employed in LexicMap benchmarking experiments [10], and 100 plasmids from EBI database (**Tab. S1**), were searched across the ATB-HQ collection using the following LexicMap command:

> **lexicmap search -j 20 -d {lexicmap_index} {query} -o {output_alignments} --debug**

Search time was measured using Galitime (**Tab. S1**).

Phylign-Fulgor searches were performed setting its calibrated threshold with respect to LexicMap (0.007%) and the following parameters: *nb_best_hits*: 2000000; *minimap_preset*: asm20; *threads*: 20; *max_ram_gb*: 300.

## SUPPLEMENTARY NOTES

### Note S1. Biological ques6on transla6on into a *k*-mer based problem

Given a biological question of interest, we assume it can be answered using *k*-mers, and assume that it can be formalized using a *k*-mer based pseudo-algorithm involving a step of *k*-mer matching. Its specific formalization may call for different *matching strategies*, i.e. a specific combination of these three elements: *matching mode*, characteristics of the reference collection, and query type.

As an example, we describe two questions recently discussed in literature (**Fig 1a**), highlighting the role of *k-*mer matching and the specific matching strategies employed for addressing the problem.

The first example was provided by Bradley and colleagues [17], who used *k*-mers to characterize plasmid host-range in reference collection. Their bioinformatic method involved an initial step of *k*-mer matching finalized to the detection of putative conjugative systems by searching for exact matches between the alleles of two specific types of conjugative genes, namely MOB and T4SS, and the sequences in the reference collection. Post-processing analysis followed *k-*mer matching to determine the presence of these putative conjugative systems, defined by the presence of at least one allele for both types of genes.

The second example was provided by Břinda and colleagues [8], who developed the Genomic Neighbour Typing (GNT) technique for predicting the phenotypic antimicrobial resistance profile of bacterial isolates. In this case, the step of *k*-mer matching is performed by ProPhyle whose role is to identify the “nearest neighbors” of the bacterial isolate of interest in a reference collection. The matching strategy used in GNT was to search the set of best matching sequences for each sequencing long-reads of the bacterial isolate. Post-processing analysis followed *k*-mer matching to merge the results of each reads and to predict the isolate’s AMR phenotypic profile based on the phenotypic profile of its nearest neighbors.

### Note S2. *k*-mer index selec6on

Selection of the most suitable *k*-mer index for a given application should be driven first by its compatibility with both the subindex paradigm and the defined matching strategy, and then by its compression and searching performances. Ideally, these evaluations should allow users to identify the best *k*-mer index that best minimizes both the search time of *k*-mer matching and the use of computational resources. If a single optimal solution for both qualities is not available among the candidates, a Pareto optimization analysis should be performed to decide whether to choose the best trade-off index or a combination of multiple *k*-mer index types. To validate the final choice (either a single or multiple indexes combined together), the chosen *k*-mer index(es) has to be integrated in the overall bioinformatic workflow and tested as well.

### Note S3. Supported matching modes

Both Metagraph and COBS support all matching modes through their threshold-based methods (defined as *exact k-mer matching* and *approximate pattern matching* by their respective authors), where thresholds are computed as a fraction of matched *k*-mers over the total number of query *k*-mers. Specifically for the *all-best-matches* and *all-best-matches-above-threshold* modes, an additional post-processing step is required to filter the output matches and retain only the best matches. This is possible, as both indexes report the list of query-match pairs together with the corresponding number of shared *k*-mers.

In contrast, Themisto and Fulgor partially support the *all-matches-above-threshold* mode, as they both implement threshold-based searches, defined as *threshold* and *threshold-union* methods by their respective authors. In both tools, similarity can be computed in one of two ways: either considering the full set of query *k*-mers, or only the positive subset, namely those query *k-*mers that have been found at least once in the index. To ensure compatibility with the *subindex paradigm*, it is essential to choose the former option (i.e. using all query *k*-mers). Themisto supports this via the parameter “*--include-unknown-kmers”*, while Fulgor requires further minor modifications in its code to enable this option. However, this mode is only partially supported because the output does not report how many *k*-mers the query shares with each match, making it impossible to rank matches based on their “similarity” with the query.

The remaining matching modes are not supported by Themisto and Fulgor. *Exact-matches* cannot be identified because 1) they both implement the so-called *intersection* method that ignores *k*-mers that are entirely absent from the index, and 2) neither return query-match similarity information in the output. Therefore, it is not possible to distinguish true exact matches (i.e., those that contain all query *k*-mers) from matches lacking the *k*-mers absent from the index. Similarly, *All-matches-above-threshold* and *all-matches* cannot be identified by Themisto and Fulgor, as 1) neither tool implements a dedicated method for these matching modes, and 2) they do not output query–match similarity values, which prevents these modes from being derived from either intersection- or threshold-based search.

## SUPPLEMENTARY FIGURES

**Fig. S1.**
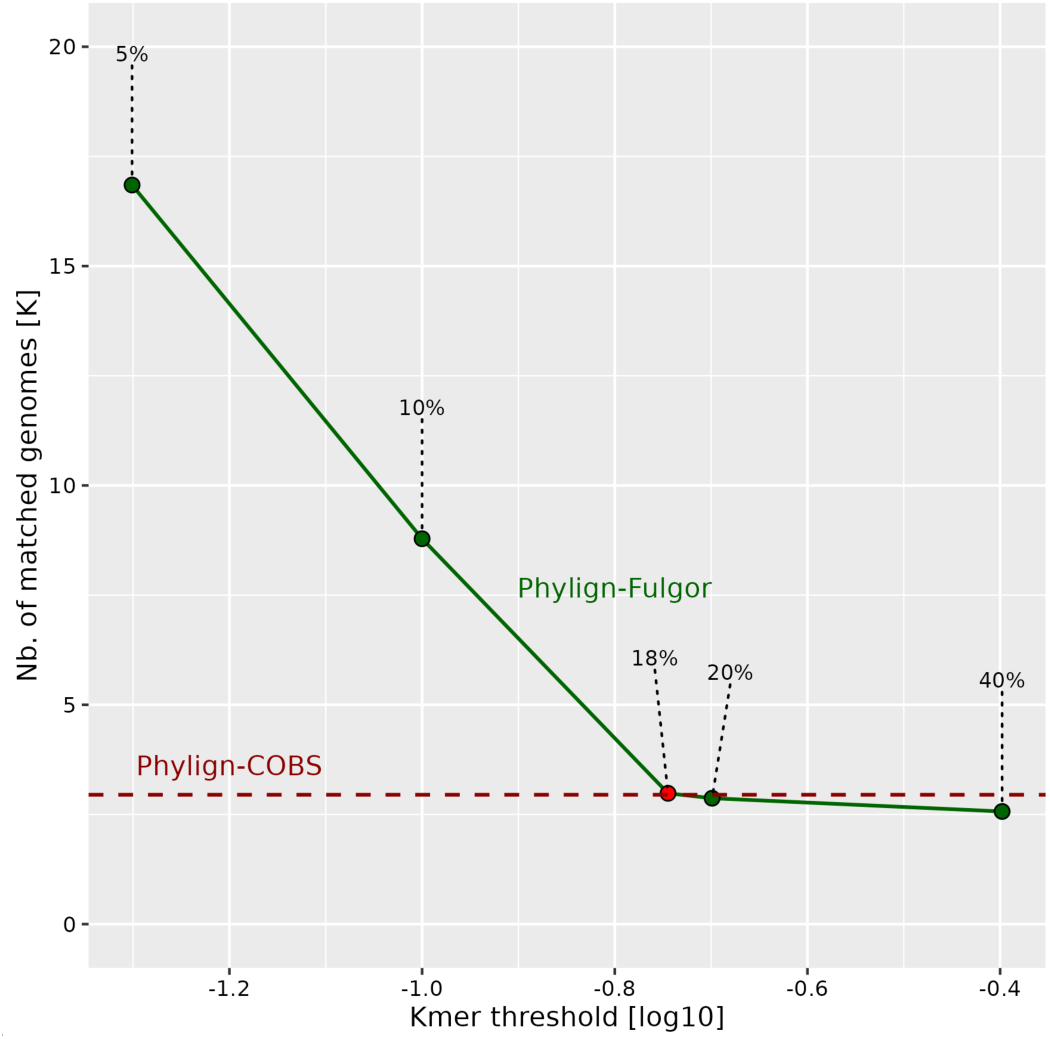
*k*-mer threshold calibration for Phylign-Fulgor’s *matches-above-threshold* using a single plasmid and 661k-HQ. The graph compares the number of matched genomes as a function of the required proportion of matching *k*-mers in the *all-matches-above-threshold* mode, in comparison with Phylign-COBS. The red point (t=18%) corresponds to the threshold yielding a similar number of matches to Phylign-COBS.

## SUPPLEMENTARY TABLES

**Tab. S1.**
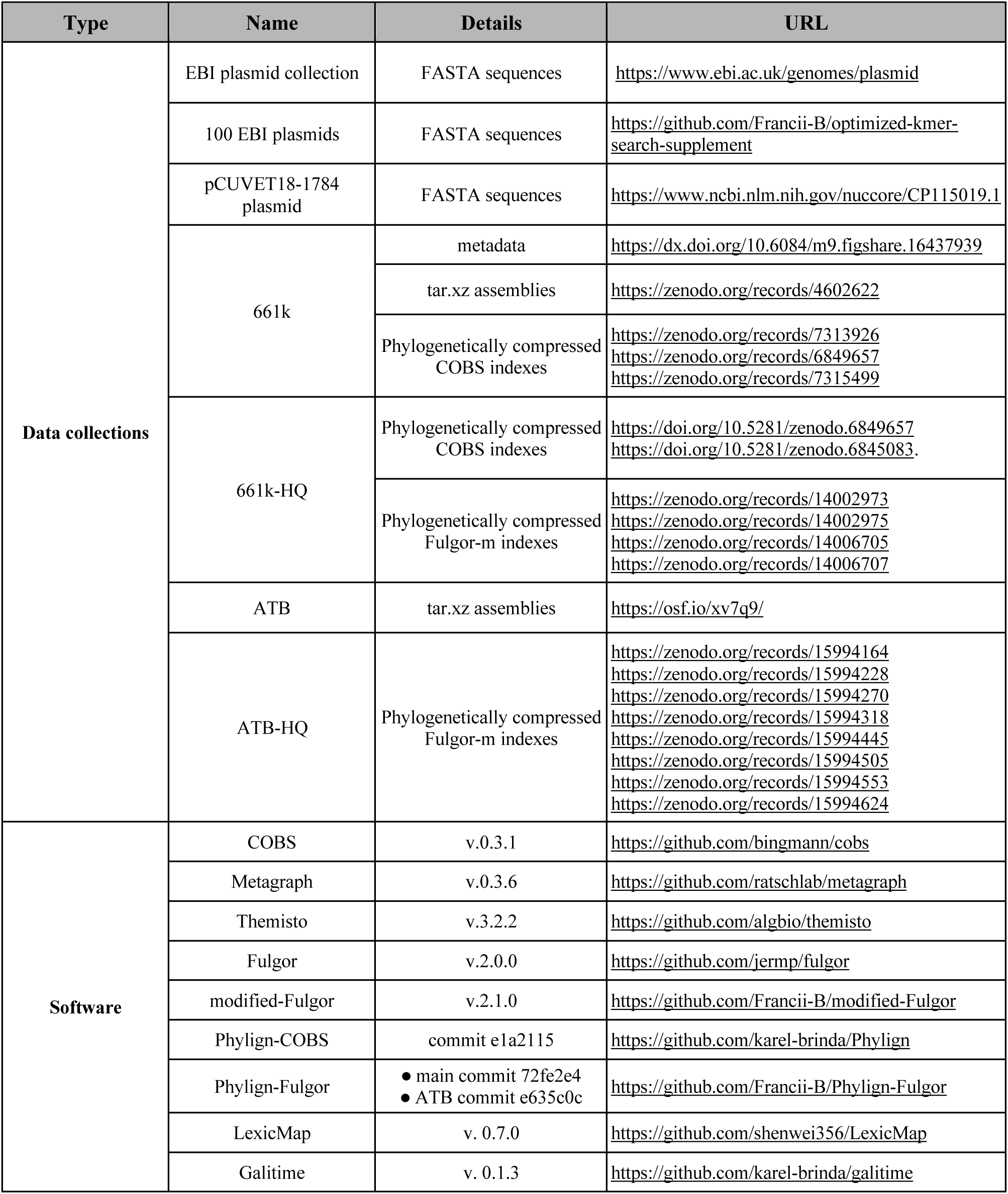
Overview of the software and data used throughout the paper.

**Tab. S2.** Detailed statitstics for BLAST-like alignments by Phylign-COBS and Phylign-Fulgor. Search time, memory requirements and outputs of the pipelines for searching all EBI plasmids (n=2,826) across the 661k-HQ collection. Both Phylign-COBS and Phylign-Fulgor were run on a MacBook Pro 18,3, setting the following parameters: *kmer_thres: 0.4; nb_best_hits: 1000; minimap_preset: asm20; threads: 4*.

| Method |  | Computational time [h] |  | Matching statistics |  |  |  |
| --- | --- | --- | --- | --- | --- | --- | --- |
|  |  | Real time | CPU time | #distinct query-genome pairs | #matched queries | #target genomes | #target batches |
| Phylign-Fulgor | match | 2.13 | 8.43 | 3,732,698 | 1,740 | 330,683 | 295 |
|  | align | 36.54 | 13.61 | 716,790 | 1,738 | 193,568 | 291 |
| Phylign-COBS | match | 3.44 | 34.97 | 9,163,123 | 1,873 | 367,565 | 298 |
|  | align | 13.20 | 14.92 | 838,830 | 1,871 | 205,231 | 296 |

**Tab. S3.** Alignment comparisons between Phylign-Fulgor and Phylign-COBS. Search time, memory usage, and the number of hits obtained when searching the pCUVET18-1784 plasmid sequence (length: 52,830 bp) across the 661k-HQ dataset (n. genomes=639,981, n. batches=305) are reported for both tools. Phylign-Fulgor and Phylign-COBS were run on the GenOuest cluster (https://www.genouest.org), using a single node with an SSD disk, 20 CPUs, and 80 GB of RAM available. The following parameters were used for Phylign-Fulgor: *nb_best_hits: 2,000,000; minimap_preset: asm20; threads: 20; max_ram_gb: 80*.

| Method |  | Matching statistics |  | Server (20 cores) |  |  |
| --- | --- | --- | --- | --- | --- | --- |
|  |  | #target genomes | #alignments | Real time [s] | CPU time [s] | RAM [GB] |
| **Single plasmid (pCUVET18-1784)** |  |  |  |  |  |  |
| Phylign-Fulgor | match (t=0.007%) | 136,777 | - | 115 | 230 | 0.5 |
|  | match+align(t=0.007%) | 114,775 | 414,865 | 3,467 | 57,788 | 3.5 |
|  | match+align(t=18%) | 2,982 | - | 70 | 145 | 0.2 |
|  | match+align(t=18%) | 2,982 | 26,079 | 617 | 5,803 | 2.8 |
|  | match (t=40%) | 2,568 | - | 128 | 176 | 0.6 |
|  | match+align(t=40%) | 2,568 | 22,956 | 642 | 5,245 | 2.4 |
| Phylign-COBS | match (t=40%) | 2,949 | - | 583 | 6,969 | 61.2 |
|  | match+align(t=40%) | 2,949 | 25,870 | 1,086 | 12,581 | 61.2 |

